# Privacy-preserving generative deep neural networks support clinical data sharing

**DOI:** 10.1101/159756

**Authors:** Brett K. Beaulieu-Jones, Zhiwei Steven Wu, Chris Williams, Ran Lee, Sanjeev P. Bhavnani, James Brian Byrd, Casey S. Greene

**Affiliations:** Genomics and Computational Biology Graduate Group, Perelman School of Medicine, University of Pennsylvania, Philadelphia, Pennsylvania, USA; University of Minnesota, Minneapolis, Minnesota, USA; Department of Systems Pharmacology and Translational Therapeutics, Perelman School of Medicine, University of Pennsylvania, Philadelphia, Pennsylvania, USA; Division of Cardiovascular Medicine, Department of Medicine, University of Michigan Medical School, Ann Arbor, Michigan, USA; Scripps Clinic and Research Foundation, San Diego, California, USA

## Abstract

**Background:** Data sharing accelerates scientific progress but sharing individual level data while preserving patient privacy presents a barrier.

**Methods and Results:** Using pairs of deep neural networks, we generated simulated, synthetic “participants” that closely resemble participants of the SPRINT trial. We showed that such paired networks can be trained with differential privacy, a formal privacy framework that limits the likelihood that queries of the synthetic participants’ data could identify a real a participant in the trial. Machine-learning predictors built on the synthetic population generalize to the original dataset. This finding suggests that the synthetic data can be shared with others, enabling them to perform hypothesis-generating analyses as though they had the original trial data.

**Conclusions:** Deep neural networks that generate synthetic participants facilitate secondary analyses and reproducible investigation of clinical datasets by enhancing data sharing while preserving participant privacy.

## Introduction

Sharing individual-level data from clinical studies remains challenging. The status quo often requires scientists to establish a formal collaboration and execute extensive data usage agreements before sharing such data. These requirements slow or even prevent data sharing between researchers in all but the closest collaborations. Individual-level data is critical for certain secondary data analyses (e.g. propensity score matching techniques) and subgroup analyses [1].

Even for efforts specifically designed to highlight the value of sharing data, investigators have been required to execute data use agreements. The New England Journal of Medicine recently held the Systolic Blood Pressure Trial (SPRINT) Data Analysis Challenge to examine possible benefits of clinical trial data sharing [2,3]. The SPRINT clinical trial examined the efficacy of intensive lowering of systolic blood pressure (<120 mmHg) compared with treatment to a standard systolic blood pressure goal (<140 mmHg). Intensive blood pressure lowering resulted in fewer cardiovascular events, and the trial was stopped early for benefit. Reanalysis of the Challenge data led to the development of personalized treatment scores [4] and decision support systems [5], in addition to a more specific analysis of blood pressure management in participants with chronic kidney disease [6]. The goal of these agreements is to to maintain participant privacy by prohibiting re-identification or unauthorized disclosure.

We sought to find a way to share data for initial and exploratory analyzes that does not require this data use agreement process. To do this we developed a technical solution for generating synthetic participants that were similar enough to the original trial data that both standard statistical and machine learning analyses yield effectively the same answers. Other methods aimed at performing this task generally fall into two groups: 1.) sampling methods with a quantifiable privacy risk [7], or 2.) Generative Adversarial Networks (GANs) [8], which are neural networks that can generate realistic data from complex distributions. In a GAN, two neural networks are trained against each other: one is trained to discriminate between real and synthetic data (the discriminator), and the other is trained to generate synthetic data (the generator). GANs have become a class of widely used machine learning methods and have recently been used in biology and medicine [9] and have been used to generate biomedical data [10,11]. However, using traditional GANs for this task provides no guarantee on what the synthetic data reveal about true participants. It is possible that the generator neural network could learn to create synthetic data that reveals actual participant data.

One way to avoid this scenario, in which a participant’s involvement in a trial could be revealed, is to limit the influence that any single study participant has on the paired neural networks’ training of one another. Differential privacy is a formal framework that provides tunable parameters that control the maximum possible privacy loss. That privacy loss comes in the form of the contribution any single individual to the results of analyses researchers perform. Nissim et al. [12] provide a particularly useful primer on understanding differential privacy to a non-technical audience as well as the implications of privacy loss in a legal context. Differential privacy has been adopted by the US Census Bureau for the 2020 US Census. The Census Bureau provides guidance on choosing an appropriate privacy loss [13,14]. A general background of differential privacy can be found in Dwork and Roth [15] and Abadi et al. introduced differential privacy for deep learning [16].

In this work, we introduce differential privacy to the GAN framework and evaluate the extent to which differentially private GANs could generate biomedical data that can be shared for valid reanalysis while controlling participant privacy risks. We achieve differential privacy by limiting the maximum influence of any single participant during training and then adding a small amount of random noise [16]. More detailed technical explanations of our usage of differential privacy can be found in the Online Methods section. We evaluated usefulness by: 1.) comparing variable distributions between the real and simulated data, 2.) comparing the correlation structure between variables in the real and simulated data, 3.) a blinded evaluation of individual-level data by three clinicians, and 4.) comparing predictors constructed on real vs. simulated data. The method generates realistic data by each of these evaluations.

## Methods

We used a type of GAN known as an Auxiliary Classifier Generative Adversarial Network (AC-GAN) [17] to simulate participants based on the population of the SPRINT clinical trial. We included all participants with measurements for the first twelve SPRINT visits (n=6,502), dividing them into a training set (n=6,000) and a test set (n=502). To evaluate the effect of applying differential privacy during the generation of synthetic participant data, we trained two AC-GANs using the training set: a traditional, standard AC-GAN (results termed “non-private” throughout the remainder of this manuscript) and an AC-GAN trained under differential privacy (results termed “private”). We used both GANs to simulate data that we then compared to the real SPRINT data by visualizing participant blood pressure trajectories, analyzing variable correlation structure and evaluating whether predictive models trained on synthetic data achieve similar performance ot models trained on real data. Three clinicians attempted to predict whether participants were real or synthetic and whether they were in the standard or intensive treatment group.

### Auxiliary Classifier GAN for SPRINT Clinical Trial Data

An AC-GAN (Supp. Fig. 1A) is made up of two neural networks competing with each other. Details about the neural network architectures are available in the Supplementary Online Methods. We trained the Generator (G) to take in a specified treatment arm (standard/intensive) and random noise and generate new participants that can fool the Discriminator (D). The generator takes in specified treatment arm in order to generate participants that belong to the specified arm. This labelling and additional task is the difference between an AC-GAN and a standard GAN. The generator simulated a systolic blood pressure, diastolic blood pressure and a number of medications for each synthetic patient for each of 12 SPRINT study visits. We trained the discriminator to differentiate real and simulated data from a dataset containing both groups. We repeated this process until the generator created synthetic participants that were difficult to discriminate from real ones (i.e. the accuracy of the discriminator could not improve much above ∼50%).

**Figure 1.**
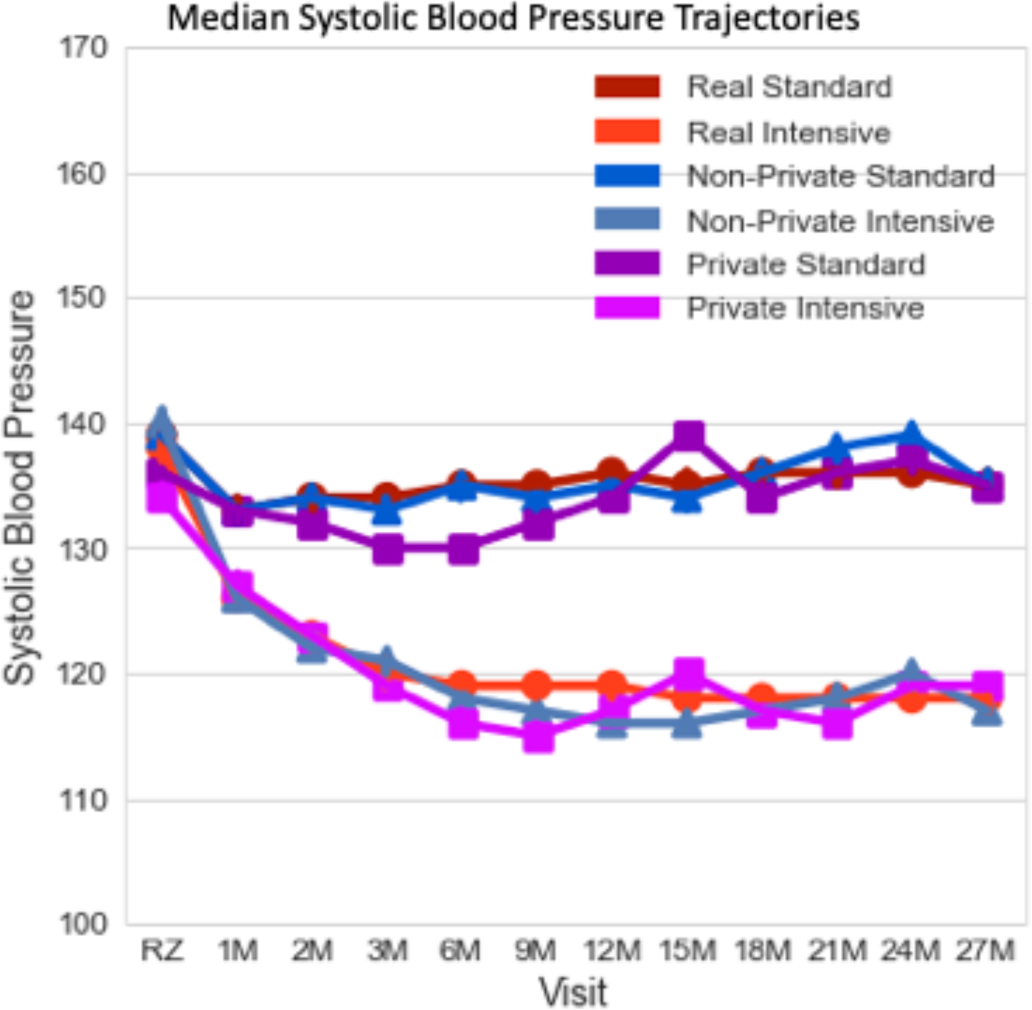
Median Systolic Blood Pressure Trajectories from initial visit to 27 months.

### Training with Differential Privacy

In order to limit the possibility that a participant’s trial involvement could be identified, we need to limit the influence any single study participant has on the neural network training of the Discriminator, the only part of the AC-GAN that accesses real data. Neural networks are trained using gradient descent, by adjusting weights according to the gradient of a loss function. Non-technically this means taking a series of steps that provide a more accurate output. To incorporate differential privacy, we limit the maximum distance of any of these steps and then add a small amount of random noise. For a detailed explanation on the processes we used please see the online methods and refer to Abadi et al. [16].

### SPRINT Clinical Trial Data

SPRINT was a randomized, single blind treatment trial that divided hypertensive participants to either intensive treatment with a systolic blood pressure target of less than 120 mmHg or standard treatment with a systolic blood pressure target of less than 140 mm Hg. The trial included a total of 9,361 participants. We included 6,502 participants who had blood pressure measurements for each of the first 12 measurements (RZ, 1M, 2M, 3M, 6M, 9M, 12M, 15M, 18M, 21M, 24M, 27M). We included measurements for systolic blood pressure, diastolic blood pressure and the count of medications prescribed to each participant, for a total of 3 parameters assessed at 12 time points.

### Clinician Evaluation

Three physicians made a blinded “real or synthetic” judgment for each of 100 figures showing systolic blood pressure, diastolic blood pressure, and number of medications at each of 12 visits. These cardiologists classified how realistic the patients looked (from 1-10 where 10 is most realistic) and whether the patients had been randomized to SPRINT’s standard or intensive treatment arm. Prior to reviewing the figures and regularly during the review of figures, the clinicians reviewed the published SPRINT protocol to help contextualize the data. We performed a Mann-Whitney U test to evaluate whether the real or synthetic samples received significantly different scores and compared the accuracy of the treatment arm classifications.

### Transfer Learning Task in the SPRINT trial

Each of the 6,502 participants in our analytical dataset was labeled by treatment arm. We evaluated machine learning methods (logistic regression, support vector machines, and random forests from the scikit-learn [33] package) by their ability to predict a participant’s treatment arm. This was done by splitting the 6,502 participants into a training set of 6,000 participants (referred to as ‘real’ in this manuscript) and a test set of 502 participants. We then trained two AC-GANs using the 6,000-participant training set, 1.) an AC-GAN model trained without differential privacy (referred to as ‘non-private’) and 2.) an AC-GAN trained with differential privacy (referred to as ‘private’). Each classifier was then trained on three datasets, 1.) the real training dataset, 2.) synthetic participants generated by the non-private AC-GAN and 3.) synthetic participants generated by the private AC-GAN. Each classifier was then evaluated on the same, real test set of participants. This allows for a comparison of classification performance between models trained on the real data, synthetic data and private synthetic data. We evaluated both accuracy as well as the correlation between important features (random forest) and model coefficients (logistic regression and support vector machine).

### Predicting Heart Failure in the MIMIC Critical Care Database

We generated synthetic patients for the purpose of predicting heart failure. MIMIC is a database of 46,297 de-identified electronic health records for critical care patients at Beth Israel. We defined patients who suffered from heart failure as any patient in MIMIC diagnosed with an ICD-9 code included in the Veterans Affairs’ Chronic Heart Failure Quality Enhancement Research Initiative’s guidelines: (402.01, 402.11, 402.91, 404.01, 404.03, 404.11, 404.13, 404.91, 404.93, 428, 281.1, 428.20, 428.21, 428.22, 428.23, 428.30, 428.31, 428.32, 428.33, 428.40, 428.41, 428.42, 428.43, and 428.9). We performed complete case analysis for patients with at least five measurements for mean arterial blood pressure, arterial systolic and diastolic blood pressures, beats per minute, respiration rate, peripheral capillary oxygen saturation (SpO2), mean non-invasive blood pressure and mean systolic and diastolic blood pressures. For patients with more than five measurements for these values, the first five were used. This yielded 8,260 total patients and 2,110 cases of heart failure. We included the first 7,500 patients in the training set and the remaining 760 in a validation set. The training and transfer learning procedures matched the SPRINT protocol.

## Results

We trained a differentially private AC-GAN to generate synthetic participants that resemble the real trial participants (Figure 1). We compare the median systolic blood pressures over time (Figure 2) of three groups, 1.) real participants (“real”), 2.) simulated participants via a non-private AC-GAN (“non-private”) and 3.) simulated participants via the differentially private AC-GAN (“private”). The non-private participants generated at the end of training appear similar to the real participants. The private participants have wider variability because of the noise added during training (Fig. 1A).

**Fig. 2.**
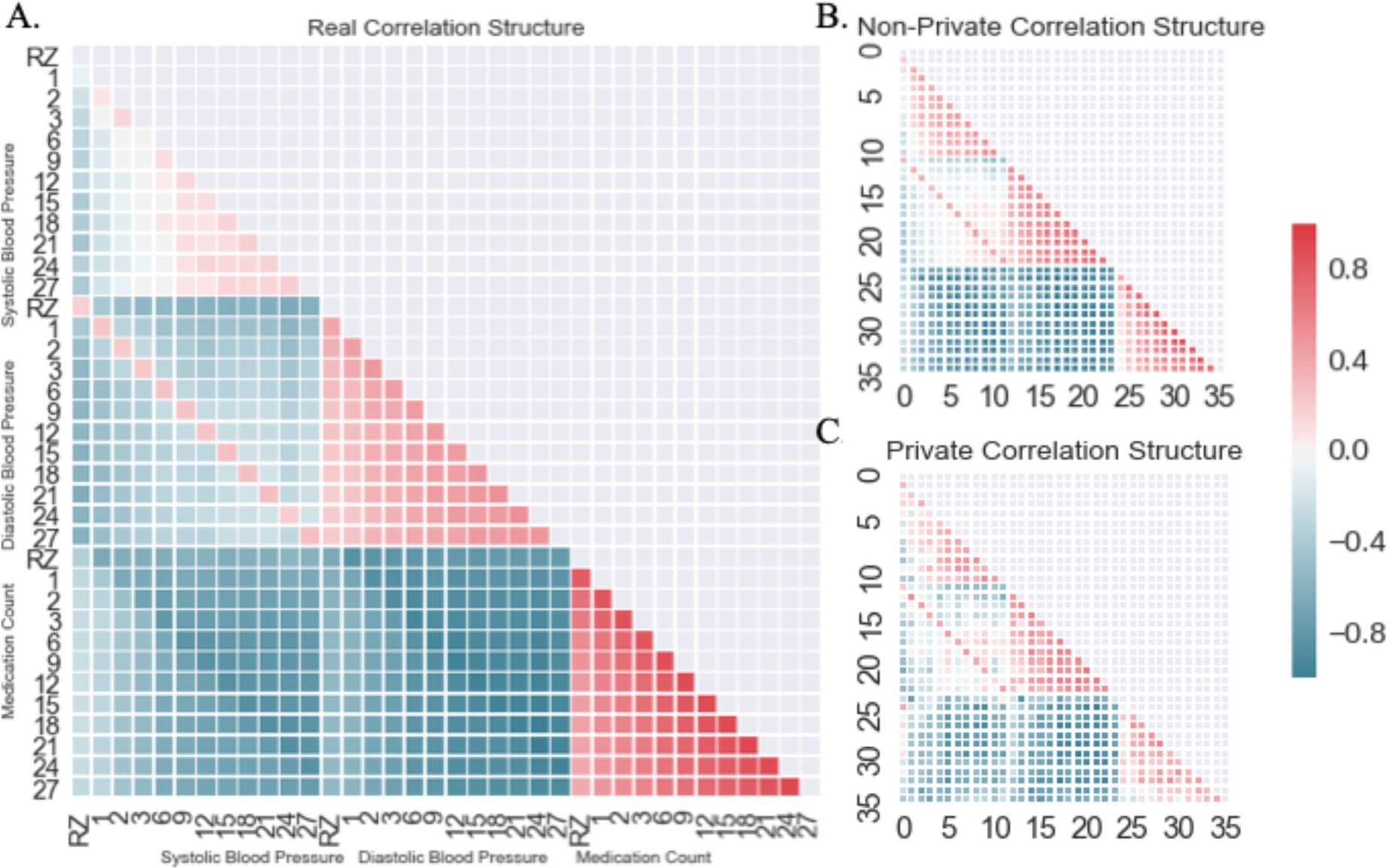
Pairwise Pearson correlation between columns for the **A.)** Original, real data, **B.)** Non-private, AC-GAN simulated data **C.)** Differentially private, AC-GAN simulated data. (RZ, randomization visit; 1M, 1 month visit; 2M, 2 month visit; 3M, 3 month visit; 6M, 6 month visit; 9M, 9 month visit; 12M, 12 month visit; 15M, 15 month visit; 18M, 18 month visit; 21M, 21 month visit; 24M, 24 month visit; 27M, 27 month visit).

Table 1 compares how close statistics calculated between the three groups were, as well as a comparison of treatment decisions between the real and synthetic participants. In particular, we examined the proportion of times an additional medication was added when a participant was above the target systolic blood pressure goal for their treatment arm (120 mm Hg for intensive, 140 mm Hg for standard). For this task the private synthetic participants closely reflected the original trial (15.51% vs 15.14%). This demonstrates the potential to meaningfully ask questions using synthetic data prior to acquiring and confirming a putative relationship in the real data.

**Table 1.** Summary statistic comparison between Real, Non-Private Synthetic and Private Synthetic Participants - mean (standard deviation).

|  | Real | Non-Private Synthetic | Private Synthetic |
| --- | --- | --- | --- |
| Systolic Blood Pressure | 129.01 (15.14) | 128.96 (14.76) | 128.74 (15.21) |
| Diastolic Blood Pressure | 72.02 (11.43) | 72.86 (10.93) | 72.92 (11.47) |
| Medications | 2.27 (1.15) | 2.04 (1.12) | 2.25 (1.14) |
| % SBP above target | 39.56% | 40.48% | 39.80% |
| % SBP above target where medication added | 15.14% | 14.99% | 15.51% |

As another method of determining whether the resulting synthetic data are similar to the real data, we measured the correlation between each study visit’s systolic blood pressure, diastolic blood pressure, and medication count. We performed this analysis within the SPRINT dataset (“real correlation structure”) and within the datasets generated by the GAN without and the GAN with differential privacy (“non-private correlation structure” and “private correlation structure,” respectively). The Pearson correlation structure of the real SPRINT data (Fig. 2A) was closely reflected by the correlation structure of the non-private generated data (Fig. 2B). Of note was initial positive correlation between the number of medications a participant was taking and the early systolic blood pressures, but this correlation decreased as time goes on. The correlation matrices between the real SPRINT data (i.e., the training data) and the non-private data were highly correlated (Spearman correlation = 0.9645, p-value < 0.0001). Addition of differential privacy during the synthetic data generation process (i.e., the “private dataset”) generated data generally reflecting these trends, but with an increased level of noise (Fig. 2C). The correlation matrices between the real SPRINT data and the private generated data were only slightly less correlated (Spearman correlation = 0.9185, p-value < 0.0001). The noisy training process of the private discriminator places an upper bound on its ability to fit the distribution of data. Increased sample sizes (such as in EHRs or other real-world data sources) would help to clarify this distribution and because larger sample sizes cause less privacy loss, less noise would need to be added to achieve an acceptable privacy budget.

### Human Comparison of Real vs. Synthetic Participants

To ensure similarity between the synthetic and real SPRINT data persists during rigorous inspection at more granular scale, we asked three clinicians to judge whether individual participant data were real SPRINT data, or synthetic data. These three physicians, experienced in the treatment of hypertension and familiar with the SPRINT trial, were asked to determine in a blinded fashion whether 100 participants (50 real, 50 synthetic) looked real. The clinicians looked for data inconsistent with the SPRINT protocol or that otherwise appeared anomalous. For example, the clinicians were alert for instances in which the systolic blood pressure was less than 100 mm Hg, but the participant was prescribed an additional medication. The clinicians classified each record on a zero to ten realism scale (10 was the most realistic), as well as whether the data correspond to standard or intensive treatment (Fig. 3A-D). The mean realism score for synthetic patients was 5.18 and the mean score for the real patients 5.26 (Figure 3E). We performed a Mann-Whitney U test to evaluate whether the scores were drawn from significantly different distributions and found a p-value of 0.333. The clinicians correctly classified 76.7% of the real SPRINT participants and 82.7% of the synthetic participants as the standard or intensive group.

**Fig 3.**
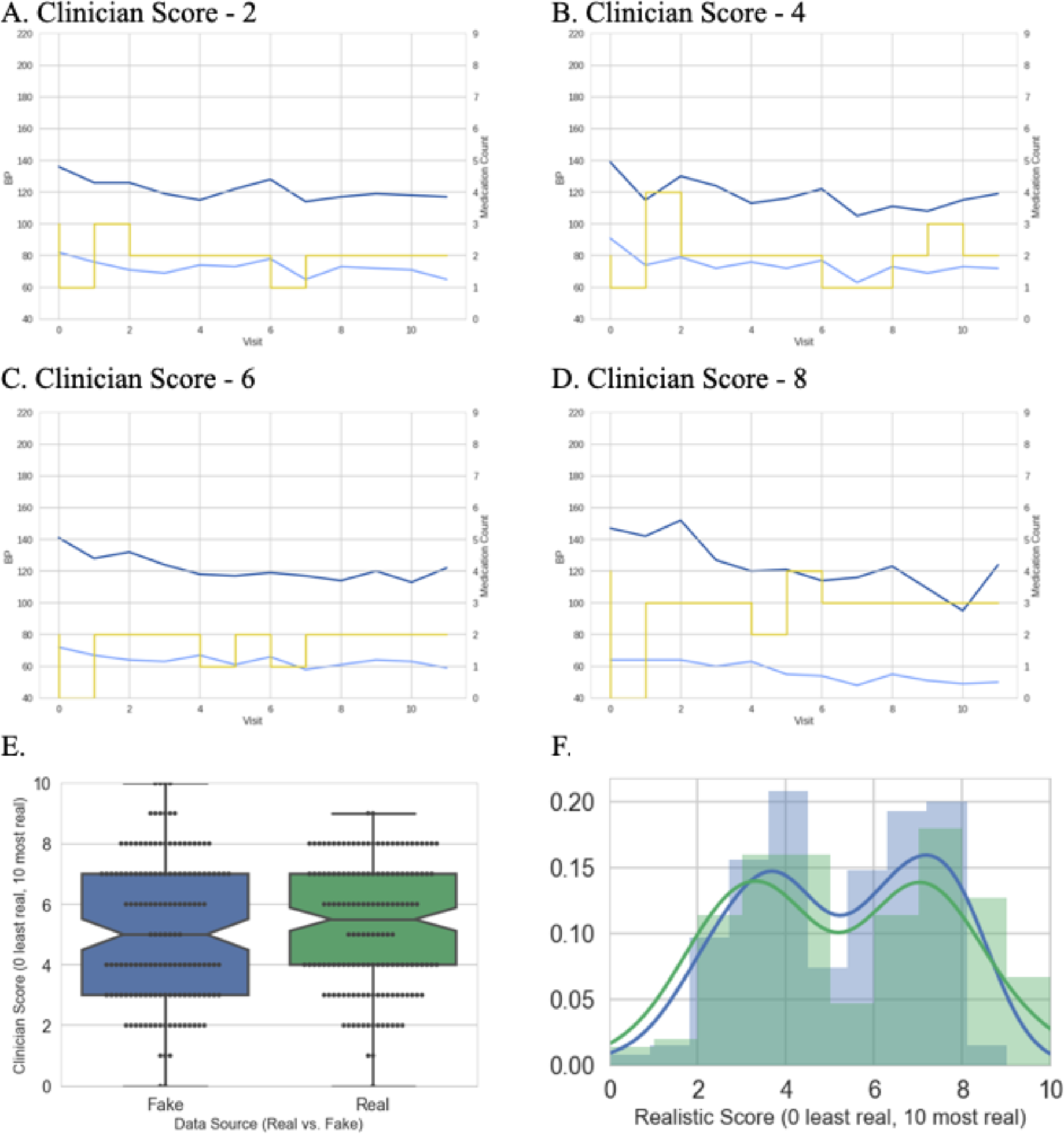
**A.)** Synthetic participant scored a 2 by clinician expert. **B.)** Synthetic participant scored a 4 by clinician expert. **C.)** Synthetic participant scored a 6 by clinician expert. **D.)** Synthetic participant scored a 8 by clinician expert. **E.)** Comparison of scores between real and synthetic participant (dotted red lines indicate means). **F.)** Distribution of scores between real (blue) and synthetic (green) patients.

### Machine Learning Models Trained on Simulated Participants are Accurate for Real Participants

Clinician review, visualizations of participant distributions and variable correlations showed that synthetic participants appeared similar to real participants. Next, we sought to determine whether or not subsequent data analyses using synthetic data matched that of the real data. To do this, we trained machine learning classifiers using four methods (logistic regression, random forests, support vector machines, and nearest neighbors) to distinguish treatment arms on three different sources of data: real participants, synthetic participants generated by the non-private model, and synthetic participants generated by the private model. We compared performance of these classifiers on a separate holdout test set of 502 real participants that were not included in the training process (Fig. 4). A drop in performance was expected because adding noise to maintain privacy reduces signal. If desired, training a non-private model could provide an approximate upper bound for expected performance.

We also sought to determine the extent to which the classifiers trained on real vs. synthetic data were relying on the same features to make their predictions (Supplemental Figure 6). We found that there was significant correlation between the importance scores (random forest) and coefficients (SVM and logistic regression) for the models trained on real vs. synthetic data (Supplemental Table 1). In addition, it is important to note that the models achieved their performance while relying on more than ten features at relatively even levels (Supplemental Figure 6), demonstrating the ability to capture multivariate correlations. Finally, we tested the correlation across cross validation folds within the real data to set an upper bound of expected correlation (Supplemental Figure 7).

**Fig 4.**
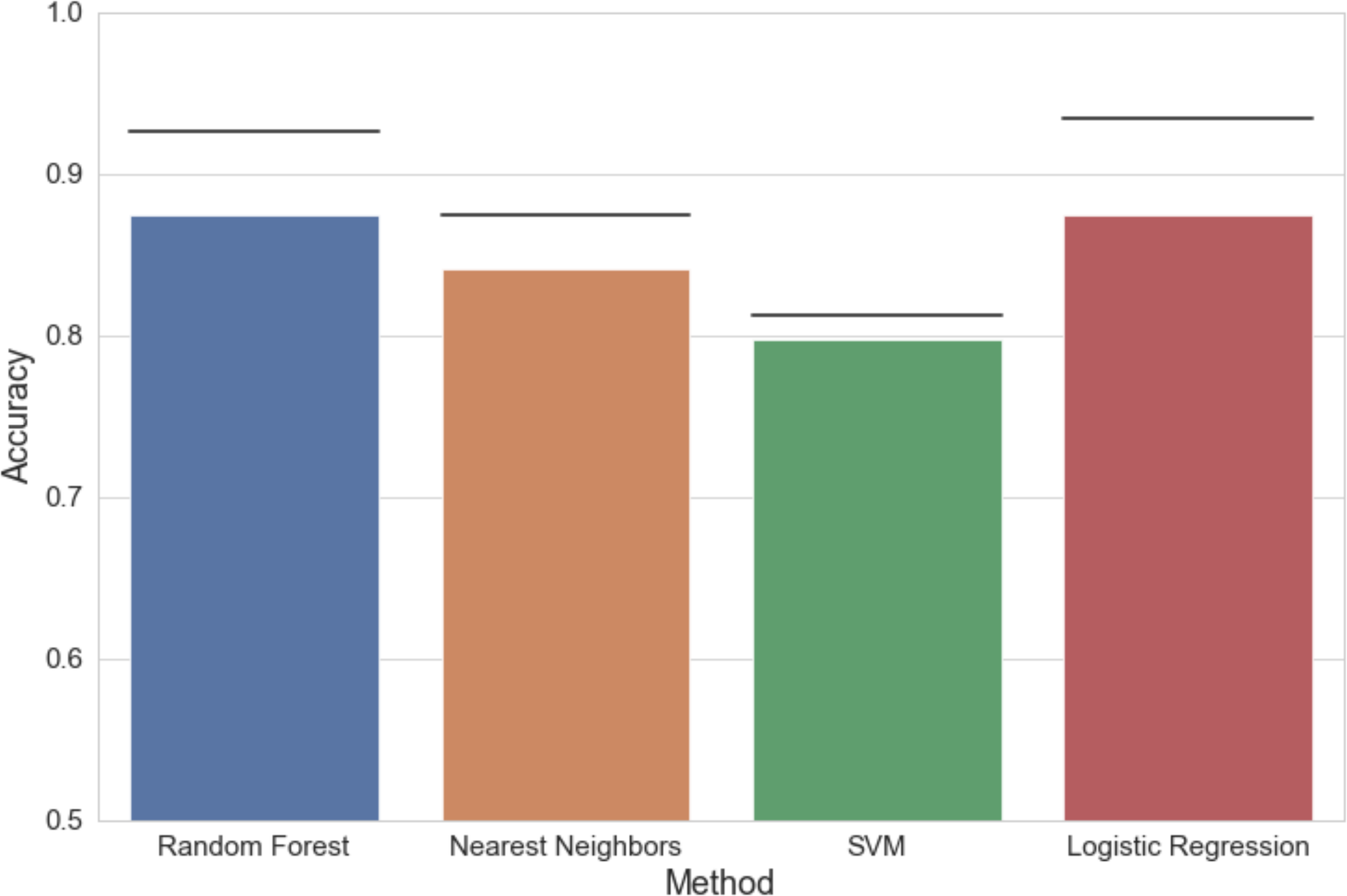
Accuracy of models trained on synthetic participants vs. real data. (Line indicates performance on real data, which on average should provide the best possible performance; bar indicates performance of classifier trained on private synthetic participants, bottom of chart indicates random performance).

**Fig 5.**
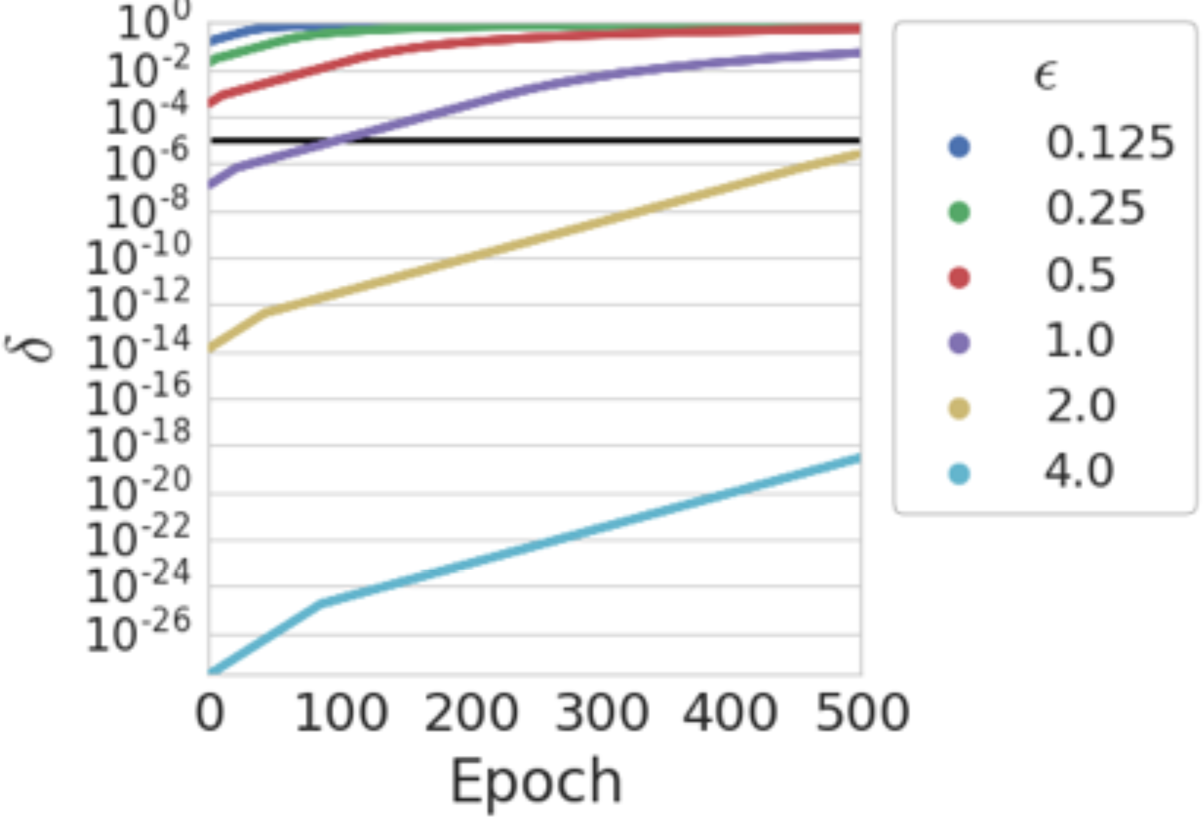
The value of delta as a function of epoch for different epsilon values. An ε value of 3.5 allows for 1000 epochs of training and δ < 10^-5^.

**Fig 6.**
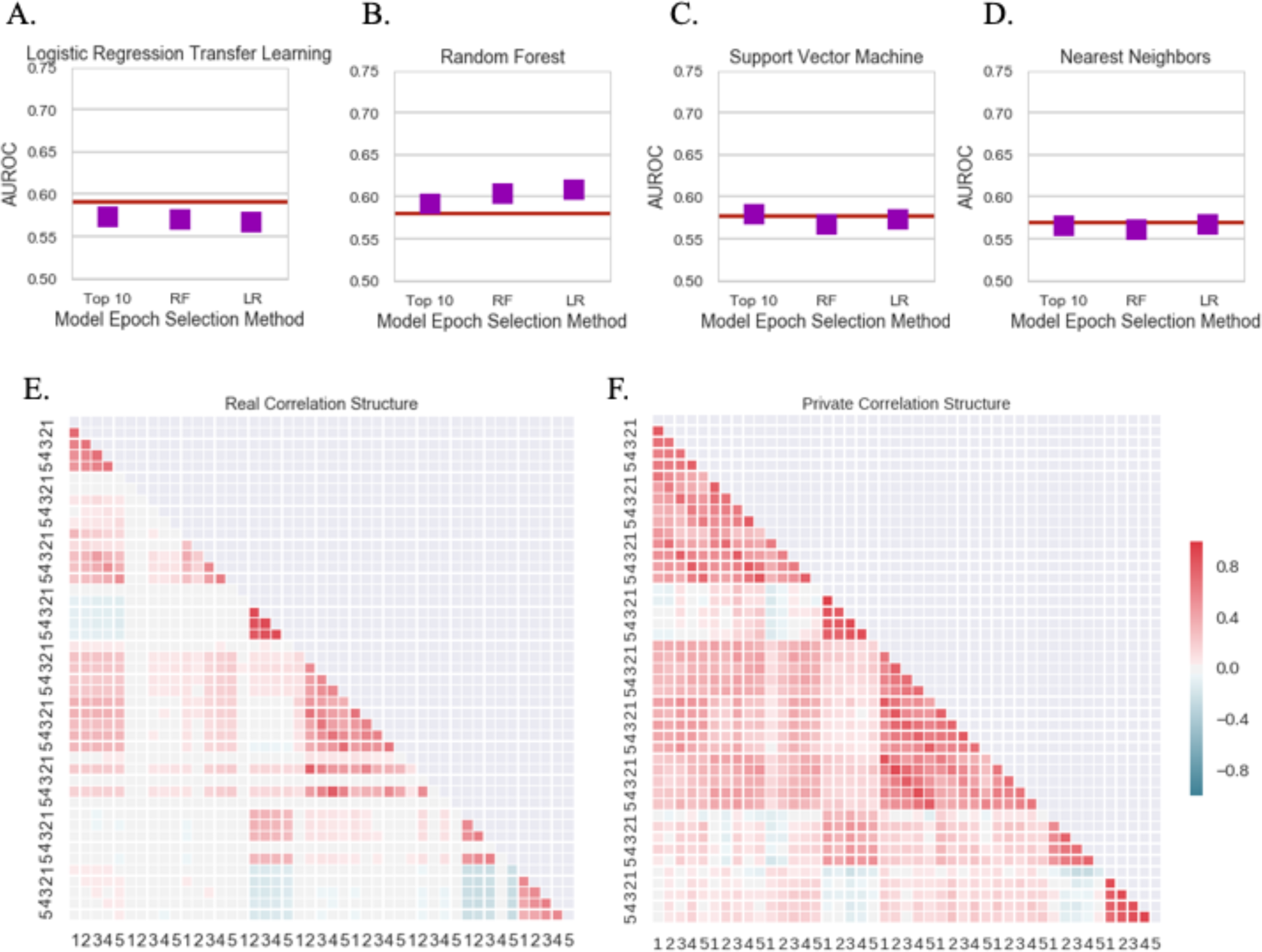
**A-D.)** Performance on transfer learning task by source of training data for each machine learning method. **E.)** Pairwise Pearson correlation between columns for the Original, real data **F.)** Pairwise Pearson correlation between columns for the Private synthetic data.

### Privacy Analysis

We evaluate privacy based on the (ε, δ) formulation of differential privacy [15]. This formal definition of differential privacy has two parameters. The parameter ε measures the maximum dataset shift that could be observed by adding or removing a single participant (termed “privacy loss”). The second parameter, δ, is the upper bound of the probability that the privacy loss exceeds ε. Put in other words, ε represents the maximum privacy loss where there is no privacy breach, and δ represents the probability of a privacy breach. We frame the problem in this way because it is impossible to anticipate all future methods of attack. For further details refer to the Online Methods section.

Therefore, it is important to choose values for ε and δ that are satisfactory to the specific use case and correspond to the consequences of a privacy breach. The values of (ε, δ) increase as the algorithm (the discriminator from the AC-GAN) accesses the private data. In our experiment, our private AC-GAN algorithm is able to generate useful synthetic data with ε = 3.5 and δ < 10^-5^ (Fig. 5). The upper bound of the epoch selection task, (see Online Methods) used (0.05, 0) per each model included for a total of (0.5, 0) differential privacy. This established a modest, single digit epsilon privacy budget of (4, 10^-5^) that is on par or lower than other methods using deep learning with differential privacy.

### Predicting Heart Failure in the MIMIC Critical Care Database

We applied the method to the MIMIC Critical Care Database [18] to demonstrate its generality. We tested whether our approach could be applied in a second dataset by predicting heart failure from the first five measurements for nine vital sign measurements in 7,222 patients. The vital sign measurements included: mean arterial blood pressure, arterial systolic and diastolic blood pressures, beats per minute, respiration rate, peripheral capillary oxygen saturation (SpO2), mean non-invasive blood pressure, and mean systolic and diastolic blood pressures. Performance on privately generated synthetic patients was on par with performance models trained on real patients (Fig. 6A-D). As in the SPRINT data, the coefficients for logistic regression and the support vector machine as well as the feature importances were significantly correlated between real and synthetic data (Supplemental Table 2).

## Discussion

Deep generative adversarial networks and differential privacy offer a technical solution to the challenge of sharing biomedical data to facilitate exploratory analyses. Our approach, which uses deep neural networks for data simulation, can generate synthetic data to be distributed and used for secondary analysis. We perform training with a differential privacy framework that limits study participants’ privacy risk. We apply this approach to data from the SPRINT clinical trial due to its recent use for a data reanalysis challenge

We introduce an approach that samples from multiple epochs to improve performance while maintaining privacy. However, this is an early stage work and several challenges remain. Deep learning models have many training parameters and require substantial sample sizes, which can hamper this method’s use for small clinical trials or targeted studies. In this work, we demonstrated the ability to use differentially private AC-GANs on relatively low-dimensional time series data sets. We applied our method to time series as we believe this provided a better test than simple point in time data because there would be time-based correlation structures. We expect this approach to be most well suited to sharing specific variables from clinical trials to enable wide sharing of data with similar properties to the actual data. We do not intend the method to be applied to generate high dimensional genetic data from whole genome sequences or other such features. Application to that problem would require the selection of a subset of variants of interest or substantial additional methodological work.

Another fruitful area of use may be large electronic health records systems, where the ability to share synthetic data may aid methods development and the initial discovery of predictive models. Similarly, financial institutions or other organizations that use outside contractors or consultants to develop risk models might choose to share generated data instead of actual client data. In very large datasets, there is evidence that differential privacy may even prevent overfitting to reduce the error of subsequent predictions.

Though our approach provides a general framing, the precise neural network architecture may need to be tuned for specific use cases. Data with multiple types presents a challenge. EHRs contain binary, categorical, ordinal and continuous data. Neural networks require these types to be encoded and normalized, a process that can reduce signal and increase the dimensionality of data. New neural networks have been designed to deal more effectively with discrete data [19,20]. Researchers will need to incorporate these techniques and develop new methods for mixed types if their use case requires it.

Due to the fluid nature of security and best practices, it is important to choose a method which is mathematically provable and ensures that any outputs are robust to post-processing. Differential privacy satisfies both needs and is thus being relied upon in the upcoming 2020 United States Census [21]. It is imperative to remember that to receive the guarantees of differential privacy a proper implementation is required. We believe testing frameworks to ensure accurate implementations are a promising direction for future work, particularly in domains with highly sensitive data. like healthcare.

The practice of generating data under differential privacy with deep neural networks offers a technical solution for those who wish to share data to the challenge of patient privacy. This technical work complements ongoing efforts to change the data sharing culture of clinical research.

## Acknowledgements

We thank Jason H. Moore (University of Pennsylvania), Aaron Roth (University of Pennsylvania), Gregory Way (University of Pennsylvania), Yoseph Barash (University of Pennsylvania), Anupama Jha (University of Pennsylvania) and Blanca Himes (University of Pennsylvania) for their helpful discussions. This Manuscript was prepared using SPRINT_POP Research Materials obtained from the NHLBI Biologic Specimen and Data Repository Information Coordinating Center and does not necessarily reflect the opinions or views of the SPRINT_POP or the NHLBI. We thank the participants of the SPRINT trial and the entire SPRINT Research Group. Funding: This work was supported by the Gordon and Betty Moore Foundation under a Data Driven Discovery Investigator Award to C.S.G. (GBMF 4552). B.K.B.-J. was supported by a Commonwealth Universal Research Enhancement (CURE) Program grant from the Pennsylvania Department of Health and by US National Institutes of Health grants AI116794, LM010098 and T15LM007092. Z.S.W is funded in part by a subcontract on the DARPA Brandeis project and a grant from the Sloan Foundation. J.B.B. is funded by US National Institutes of Health grant K23-HL128909. Author Contributions: B.K.B.-J. and C.S.G. conceived the study. B.K.B.-J. And C.W. performed initial analyses. B.K.B.-J. and Z.S.W. designed and validated the privacy approach. J.B.B performed a blinded review of records. B.K.B.-J., C.S.G. and Z.S.W. wrote the manuscript and all authors revised and approved the final manuscript. Competing interests: The authors have no competing interests to disclose. Data and materials availability: All data used in this manuscript are available via the NHLBI (https://biolincc.nhlbi.nih.gov/studies/sprint_pop/), the source code is available via GitHub (https://github.com/greenelab/SPRINT_gan) and an archived version is available via Figshare (DOI: 10.6084/m9.figshare.5165737).

## Supplementary Online Methods

We developed an approach to train auxiliary classifier generative adversarial networks (AC-GANs) in a differentially private manner to enable privacy preserving data sharing. Generative adversarial networks offer the ability to simulate realistic-looking data that closely matches the distribution of the source data.

AC-GANs add the ability to generate labeled samples. By training AC-GANs under the differential privacy framework we generated realistic samples that can be used for initial analysis while guaranteeing a specified level of participant privacy.

The source code for all analyses is available under a permissive open source license in our repository (https://github.com/greenelab/SPRINT_gan). In addition, continuous analysis [22] was used to re-run all analyses, to generate docker images matching the environment of the original analysis, and to track intermediate results and logs. These artifacts are freely available (https://hub.docker.com/r/brettbj/sprint-gan/ and archival version: https://doi.org/10.6084/m9.figshare.5165731.v1).

### Background

A pair of recent preprints have reported generation of synthetic individual participant data via neural networks [10,11]. For example, Esteban et al., generated synthetic patient data and showed that a neural network could not distinguish between the synthetic data and real data. However, it is not enough to simply build synthetic participants. Numerous linkage and membership inference attacks on both biomedical datasets [23–30] and from machine learning models [31–33] have demonstrated the ability to re-identify participants or reveal participation in a study.

To provide a formal privacy guarantee, we built GANs to generate realistic synthetic individual participant data with mathematical properties like those of the original participants’ data, adding the extra protection of differential privacy [15]. Differential privacy protects against common privacy attacks including membership inference, homogeneity and background knowledge attacks. Informally, differential privacy requires that no single study participant has a significant influence on the information released by the algorithm (see Materials and Methods for a formal definition). Despite being a stringent notion, differential privacy allows us to generate new plausible individuals while revealing almost nothing about any single study participant. Within the biomedical domain, Simmons and Berger developed a method using differential privacy to enable privacy preserving genome-wide association studies [34]. Recently, methods have also been developed to train deep neural networks under differential privacy with formal assurances about privacy risks [16,35]. In the context of a GAN, the discriminator is the only component that accesses the real, private, data. By training the discriminator under differential privacy, we can produce a differentially private GAN framework.

### Auxiliary Classifier Generative Adversarial Network

We implemented the AC-GAN as described in Odena et al. [17] using Keras [36] to simulate systolic and diastolic blood pressures as well as the number of hypertension medications prescribed. Results shown use a latent vector of dimension 100, a learning rate of 0.0002, and a batch size of 1 trained for 500 epochs. To conform with the privacy claims laid out in Abadi et al. [16], gradients must be clipped per example, in our implementation this requires the batch size to be 1. To handle edge cases and mimic the sensitivity of the real data measurements, we take the floor of zero or the simulated value and convert all values to integers. Full implementation details can be seen in the GitHub repository (https://github.com/greenelab/SPRINT_gan/blob/master/ac_gan.py).

We chose convolutional layers because of the structure imposed by sequential measurements made during the clinical trial. The features were ordered according to timing, so local structure was tied to temporality. We used deep convolutional neural networks for both the generator and discriminator (Supp. Fig. 1B, 1C).

### Differential Privacy

Differential privacy is a stability property for algorithms, specifically for randomized algorithms [37]. Informally, it requires that the change of any single data point in the data set has little influence on the output distribution by the algorithm. To formally define differential privacy, let us consider X as the set of all possible data records in our domain. A dataset is a collection of n data records from X. A pair of datasets D and D’ are neighboring if they differ by at most one data record. In the following, we will write R to denote the output range of the algorithm, which in our case correspond to the set of generative models.

#### Definition 1

[Differential Privacy [38]]: Let ε, δ > 0. An algorithm A: X^n^ → R satisfies (ε, δ)-differential privacy if for any pair of neighboring datasets D, D’, and any event S ⊆ R, the following holds

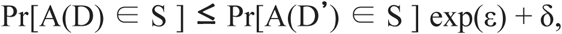

where the probability is taken over the randomness of the algorithm.

A crucial property of differential privacy is its resilience to post-processing --- any data independent post-processing procedure on the output by a private algorithm remains private.

#### More formally

Lemma [Resilience to Post-Processing]: Let algorithm A: X^n^ → R be an (ε, δ)-differentially private algorithm. Let A’: R → R’ be a “post-processing” procedure. Then their composition of running A over the dataset D, and then running A’ over the output A(D) also satisfies (ε, δ)-differential privacy.

Robustness to post-processing is critical to our application because it means that all downstream uses of the data are also (ε, δ)-differentially private. Therefore, by making the discriminator, the only part of the system that accesses the real data, differentially private, the rest of the system is also differentially private.

### Training AC-GANs in a Differentially Private Manner

We trained under differential privacy by limiting the effect any single SPRINT study participant has on the training process and by adding random noise based on the maximum effect of a single study participant. From the technical perspective, we limited the effect of participants by clipping the norm of the discriminator’s training gradient and added proportionate Gaussian noise. This combination ensures that training cannot be tied to an individual and that it could have been guided by a different subject within or outside the real training data. The maximum effect of an outlier is limited and bounded. Comparing the neural network loss functions of the private and non-private training process demonstrates the effects of these constraints. Under normal training the losses of the generator and discriminator converged to an equilibrium before eventually increasing steadily (Supp. Fig. 1D). Under differentially private training the losses converged to and remained in a noisy equilibrium (Supp. Fig. 1F). At the beginning of training the neural networks changed rapidly. As training continued and the model achieved a better fit these steps, the gradient, decreased. Eventually the gradient becomes too small in comparison to the noise for training to continue any further.

As the models achieve better fit, the gradient shrinks, causing the gradient to noise ratio to decrease. This can occasionally lead to the private generator and discriminator falling out of sync (Supp. Fig. 3) or more commonly the private model generating less realistic samples due to noise. To best select epochs, or training steps, where synthetic samples closely real samples, we tested each epoch’s data by training an additional classifier that must distinguish whether a generated participant was a part of the normal or intensive treatment groups.

During the training of AC-GAN, the only part that requires direct access to the private (real) data was the training of the discriminator. To achieve differential privacy, we only needed to “privatize” the training of the discriminators. The differential privacy guarantee of the entire AC-GAN directly followed because the output generative models are simply post-processing from the discriminator. We trained the discriminator using a differentially private version of the Adam method [39]. The standard Adam method iteratively updated the model parameters based on the gradients of the underlying loss function. To preserve privacy, we added noise to the gradient computed at each step as follows: first, we to ensured that the ℓ^2^-norm of the gradient is bounded by clipping the gradient; then we perturbed each coordinate of the gradient by adding noise drawn from the Gaussian distribution with mean 0 and standard deviation proportional to the gradient clip size. The more noise we added (relative to the clipped norm of the gradient) the better the privacy guarantee we provide.

Due to the noisy training process, the losses of the discriminator and generator do not always converge (Supp. Fig. 1) and the training algorithm may have to be rerun. To properly account the total privacy loss from all the runs, we started with a target privacy budget (given by privacy parameters ε and δ) and repeatedly ran the private training algorithm until the AC-GAN converges or the privacy budget is exhausted. We used the moments accountant described in Abadi et al. [16] to keep track of the privacy parameters (ε, δ) over time.

We clipped the ℓ^2^-norm of the gradient at 0.0001 and added noise from a normal distribution with a σ^2^ of 1 (N(µ, 1 * (0.0001^2^))). In our experiment, the AC-GAN trained in the second run of the algorithm converged, and the entire training process incurred a privacy loss within the budget (ε = 4, δ = 10^-5^).

### Differentially Private Model Selection

We trained under differential privacy by limiting the effect any single SPRINT study participant has on the training process and by adding random noise based on the maximum effect of a single study participant. To do this, we first clipped the norm of the discriminator’s training gradient. This clipping provides an upper bound on the maximum effect of a single study participant. We then added Gaussian noise according to this upper bound and the specified acceptable privacy loss. This combination ensures that training cannot be tied to an individual and that it could have been guided by a different subject within or outside the real training data. The maximum effect of an outlier is limited and bounded. Comparing the neural network loss functions of the private and non-private training process demonstrates the effects of these constraints. Under normal training the losses of the generator and discriminator converged to an equilibrium before eventually increasing steadily (Supp. Fig. 1D). Under differentially private training the losses converged to and remained in a noisy equilibrium (Supp. Fig. 1F). At the beginning of training the neural networks changed rapidly. As training continued and the model achieved a better fit these steps, the gradient, decreased. Eventually the gradient becomes too small in comparison to the noise for training to continue any further.

As the models achieve better fit, the gradient shrinks, causing the gradient to noise ratio to decrease. This can occasionally lead to the private generator and discriminator falling out of sync (Supp. Fig. 2) or more commonly the private model generating less realistic samples due to noise. To best select epochs, or training steps, where synthetic samples closely real samples, we tested each epoch’s data by training an additional classifier that must distinguish whether a generated participant was a part of the normal or intensive treatment groups.

We found that sampling from multiple different epochs throughout training provided a more diverse training set. This provided summary statistics closer to the real data and higher accuracy in the transfer learning task. During the GAN training, we saved all the generative models across all epochs. We then generated a batch of synthetic data from each generative model and used a machine learning algorithm (logistic regression or random forest) to train a prediction model based on each synthetic batch of data. We then tested each prediction model on the training set from the real dataset and calculate the resulting accuracy. To select epochs that generate training data for the most accurate models under differential privacy, we used the standard “Report Noisy Min” subroutine: first add independent Laplace noise to the accuracy of each model (drawn from Lab(1/(n*ε)) to achieve (ε, 0) differential privacy where n is the size of the private dataset we perform the prediction on and output the model with the best noisy accuracy.

In practice, we choose the top five models that performed best on the transfer learning task for the training data using both logistic regression classification and random forest classification (for a total of 10 models). We performed this task under (0.5, 0)-differential privacy. In each of the ten rounds of selection epsilon was set to 0.05. This achieves a good balance of accuracy while maintaining a reasonable privacy budget.

## Supplementary Online Results

**Supplemental Figure 1.**
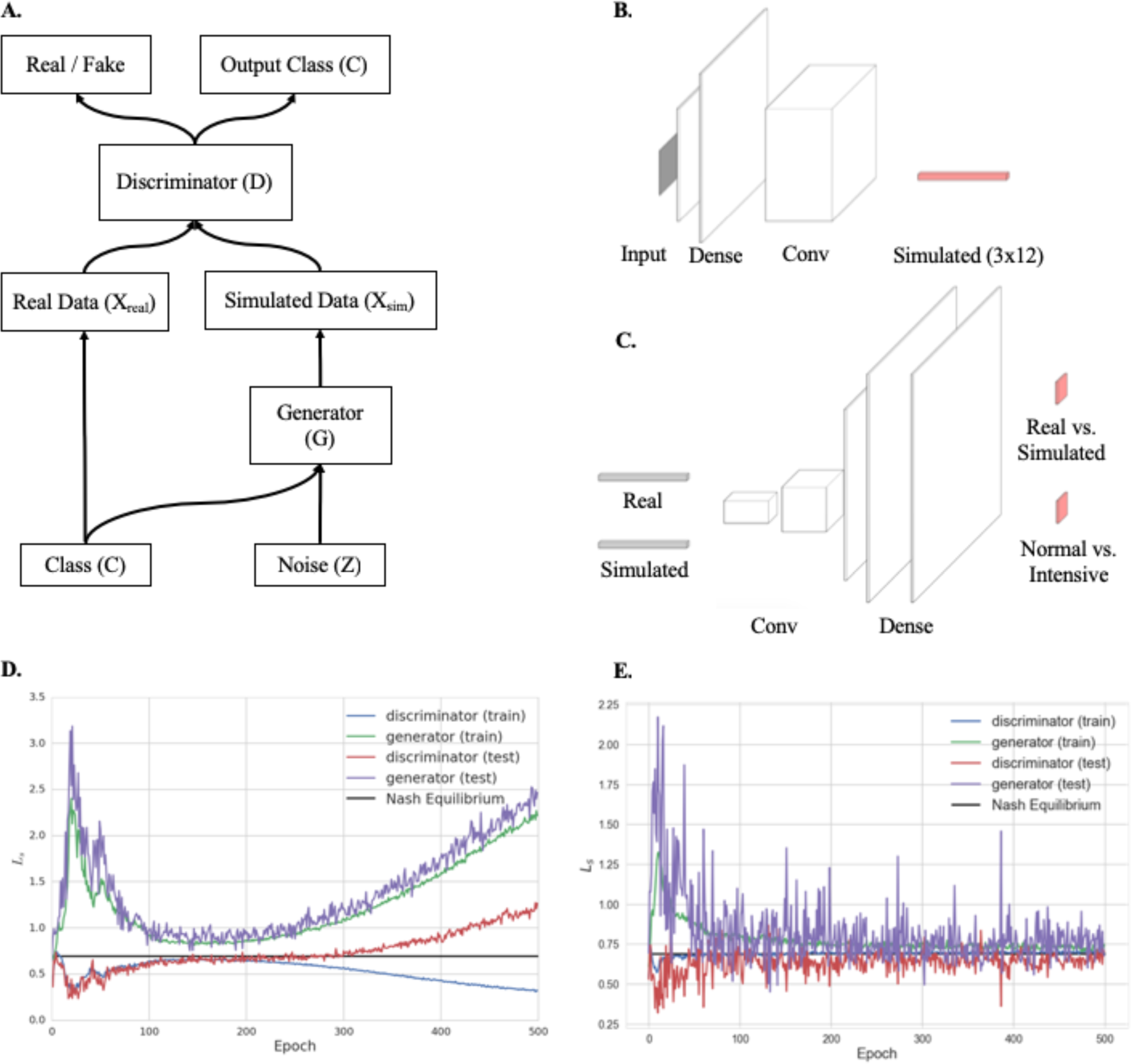
AC-GAN architecture and training. A.) Structure of an AC-GAN. B.) The generator model takes a class label representing the treatment group (e.g. intensive or standard care group) and random noise as input and outputs a 3×12 vector for each participant (SBP, DBP and medication counts at each time point). C.) The discriminator model takes both real and simulated samples as input and learns to predict the source and a class label (i.e. normal or intensive treatment group). D.) Training loss for a non-private AC-GAN. E.) Training loss for a private AC-GAN.

**Supplemental Figure 2.**
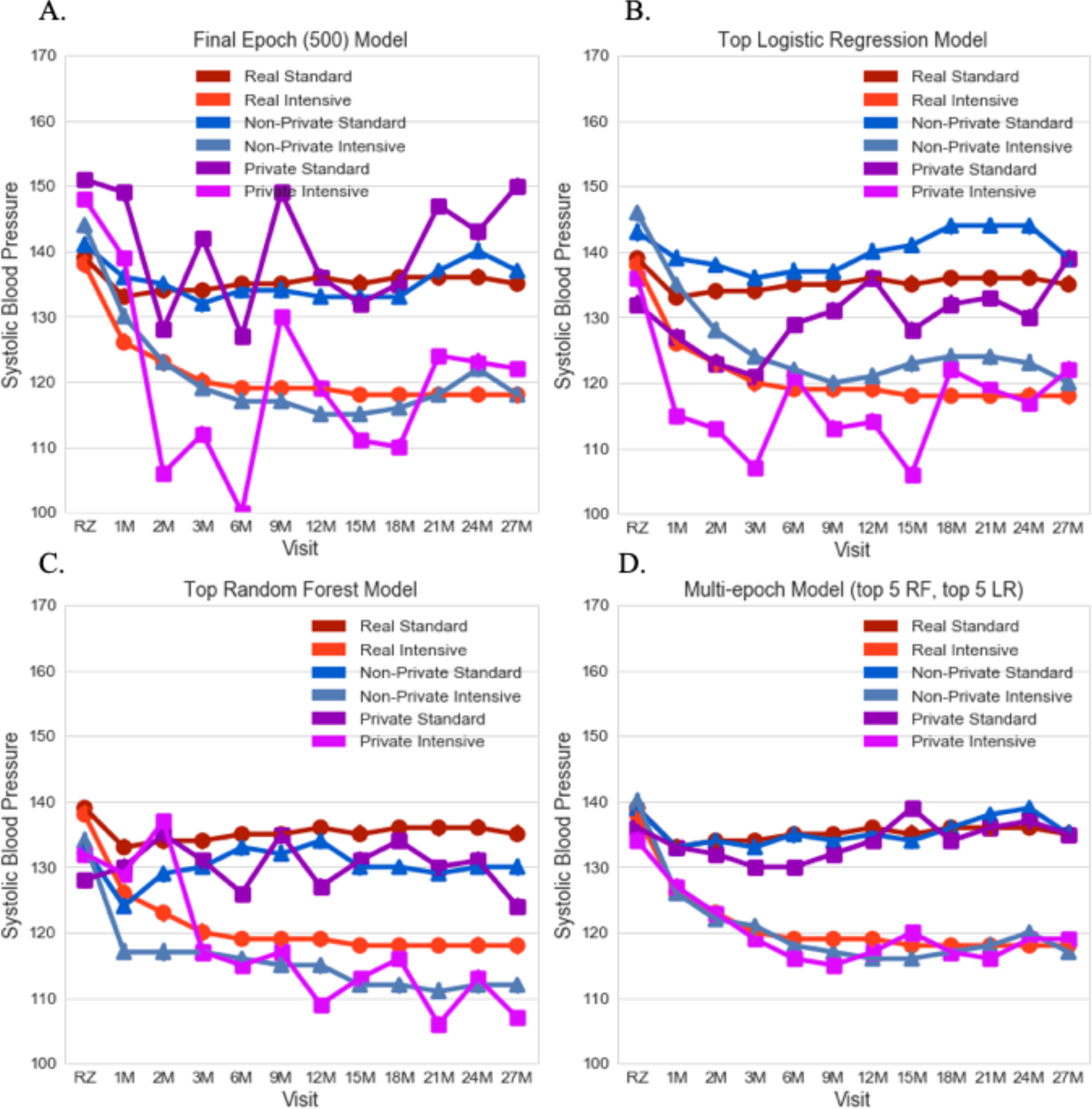
Median Systolic Blood Pressure Trajectories from initial visit to 27 months. **A.)** Simulated samples (private and non-private) generated from the final (500th) epoch of training. **B.)** Simulated samples generated from the epoch with the best performing logistic regression classifier. **C.)** Simulated samples from the epoch with the best performing random forest classifier. **D.)** Simulated samples from the top five random forest classifier epochs and top five logistic regression classifier epochs. We applied two common machine learning classification algorithms and selected the top epochs in a differentially private manner (Supp. Fig. 2B and 2C). However, selecting only a single epoch does not account for the AC-GAN training process. Because the discriminator and generator compete from epoch to epoch, their results can cycle around the underlying distribution. The non-private models consistently improved throughout training (Supp. Fig. 4A, Supp. Fig. 5A), but this could be due to the generator eventually learning characteristics specific to individual participants. We observed that epoch selection based on the training data was important for the generation of realistic populations from models that incorporated differential privacy (Supp. Fig. 4B, Supp. Fig. 5B). To address this, we simulated 1,000 participants from each of the top five epochs by both the logistic regression and random forest evaluation on the training data and combined them to form a multi-epoch training set. This process maintained differential privacy and resulted in a generated population that, throughout the trial, was consistent with the real population (Supp Fig. 2D). The epoch selection process was independent of the holdout testing data.

**Supplemental Figure 3.**
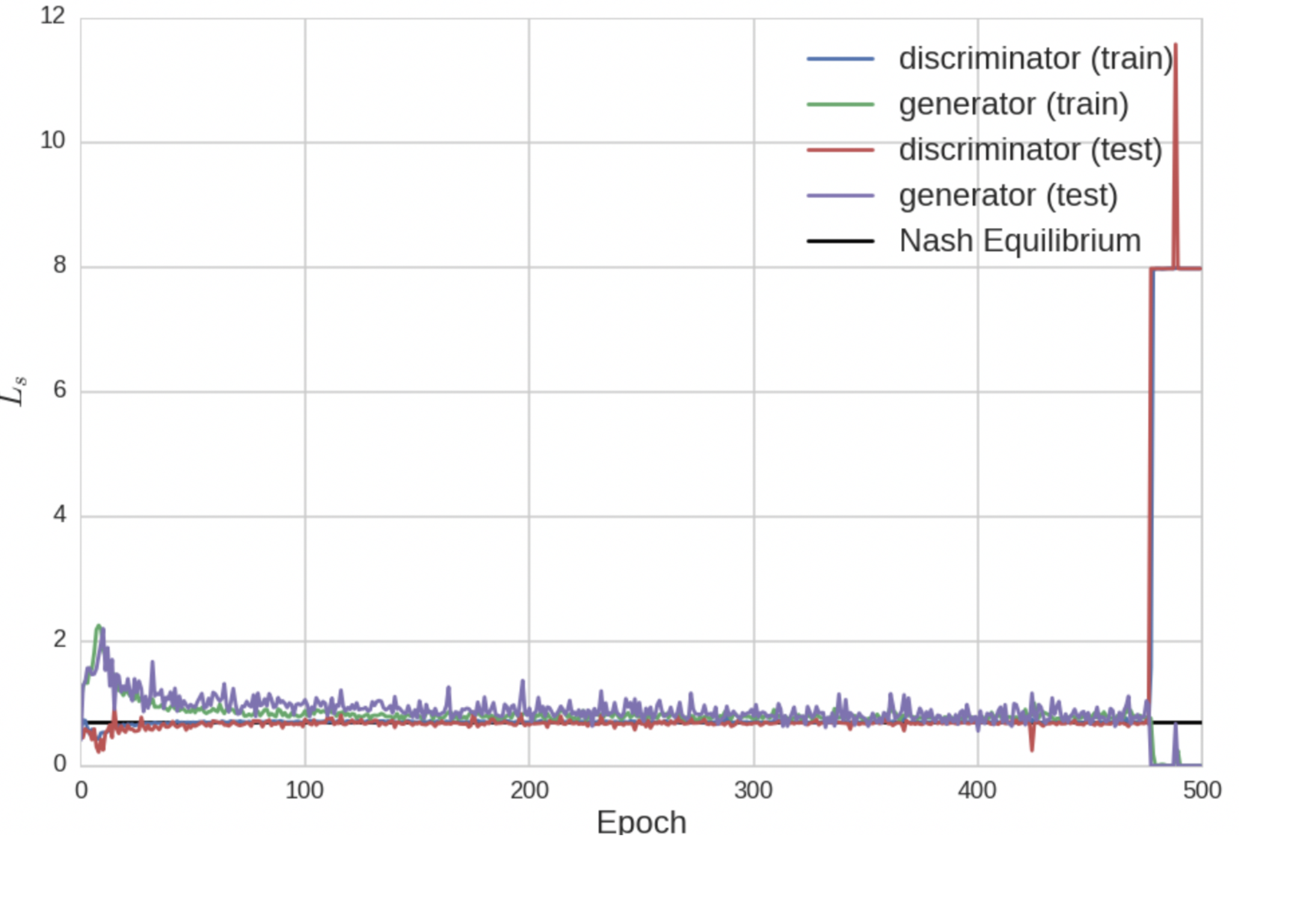
Random noise breaks equilibrium.

**Supplemental Figure 4.**
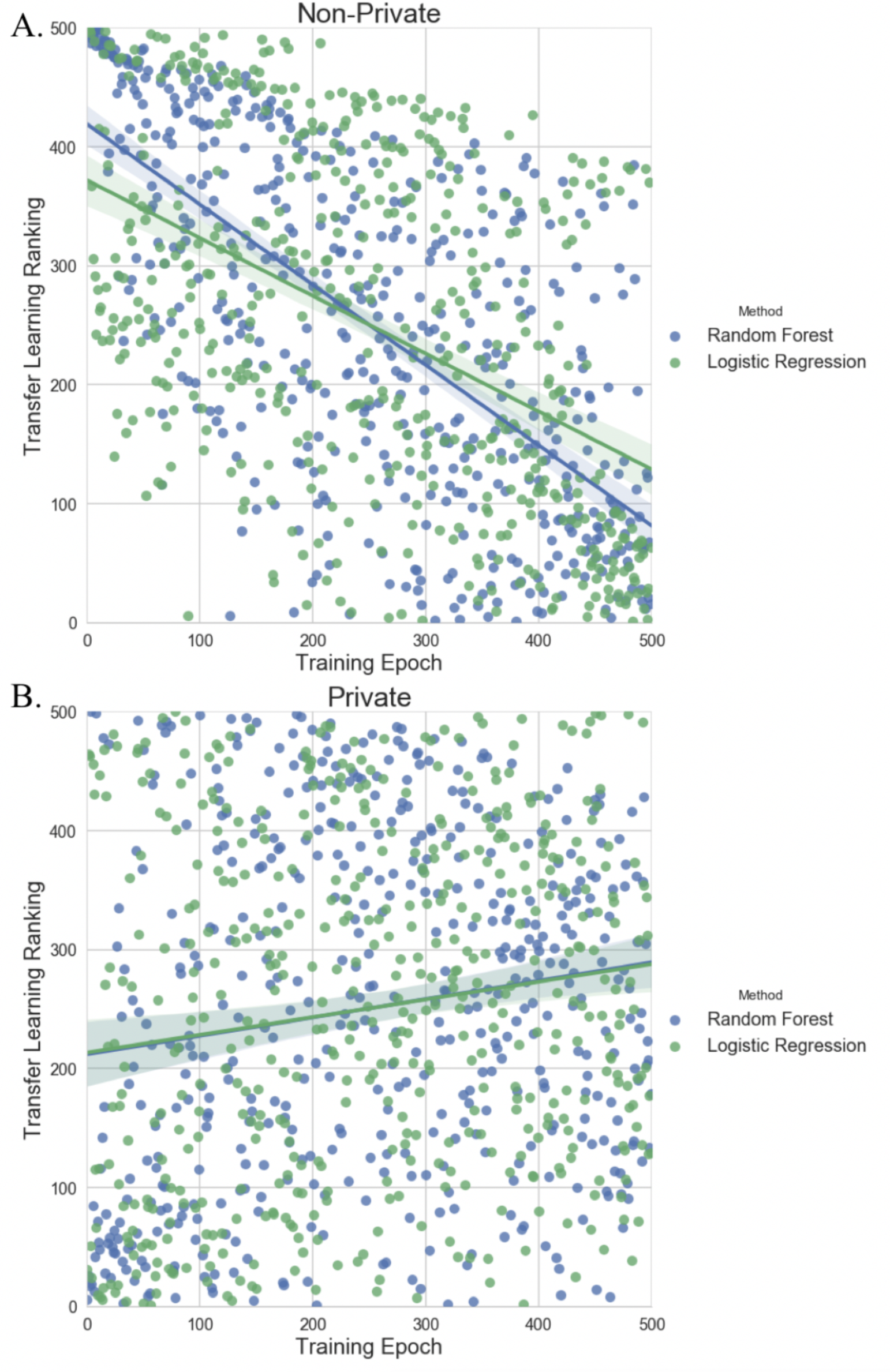
Top Ranking Epochs for Transfer Learning Exercise.

**Supplemental Figure 5.**
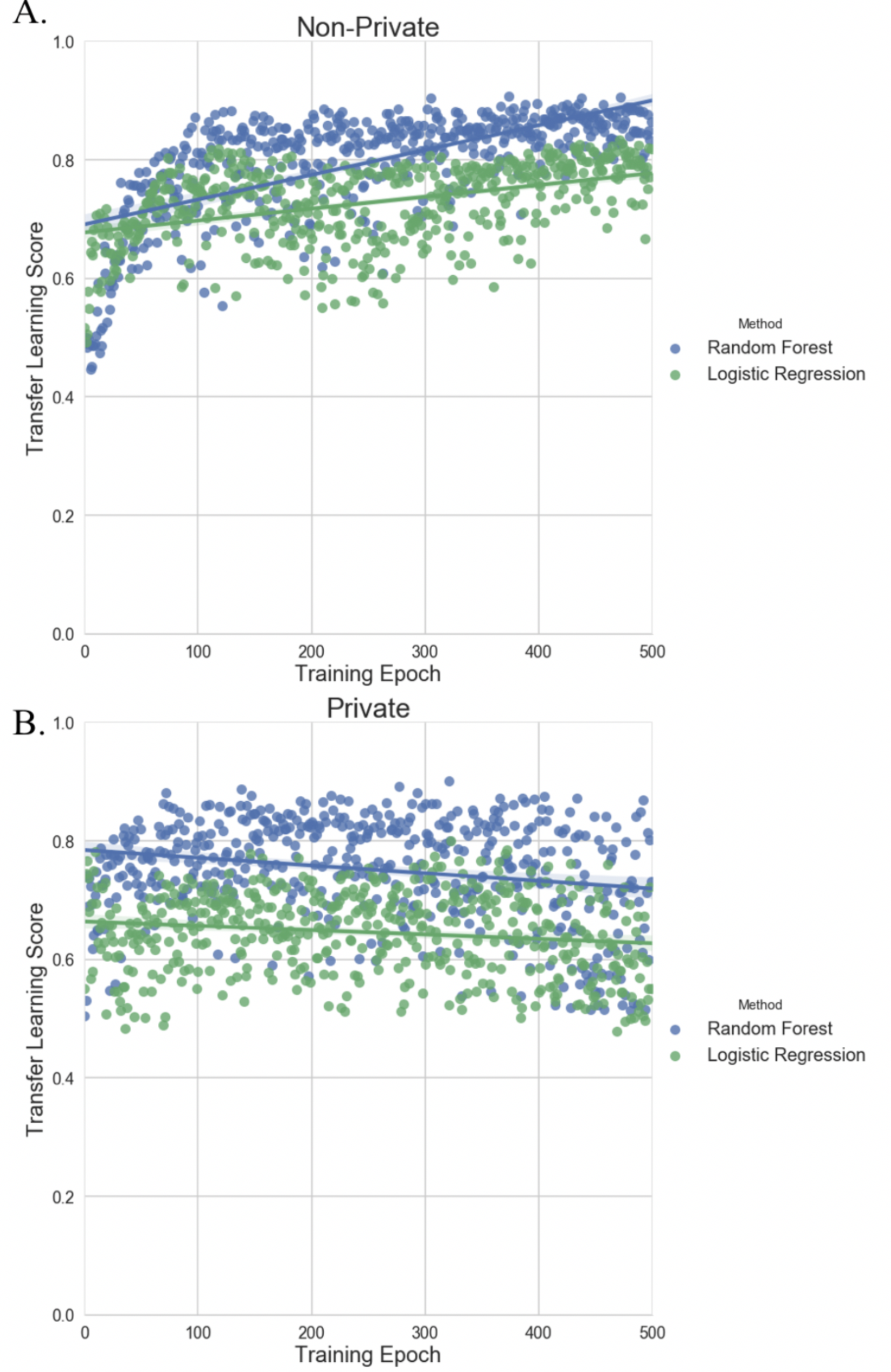
Scores vs. Epoch for Transfer Learning Task.

**Supplemental Figure 6.**
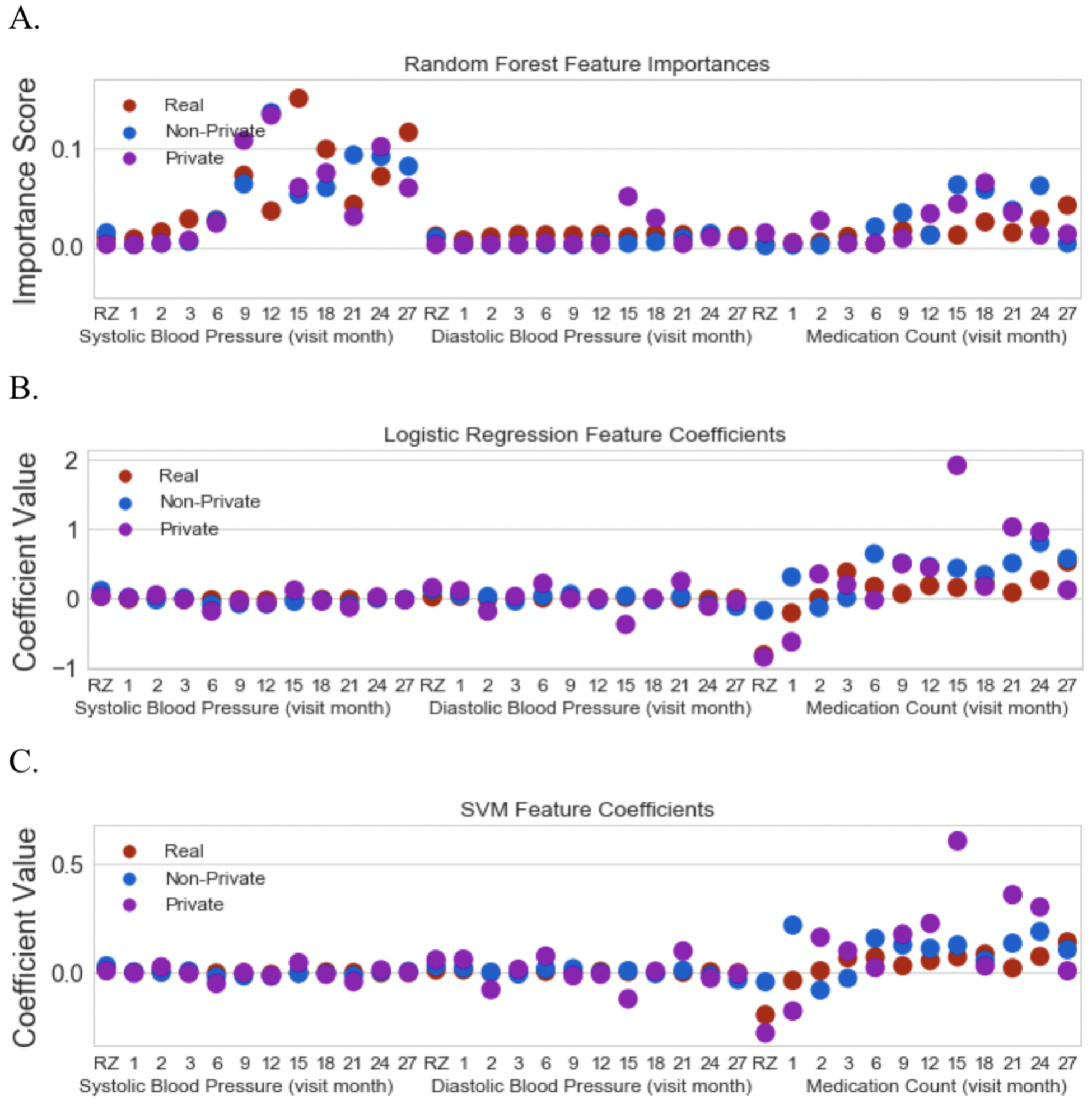
**A.)** Random forest variable importance scores by training data. **B.)** Logistic Regression variable coefficients by training data. **C.)** Support Vector Machine variable coefficients by training data.

**Supplemental Figure 7.**
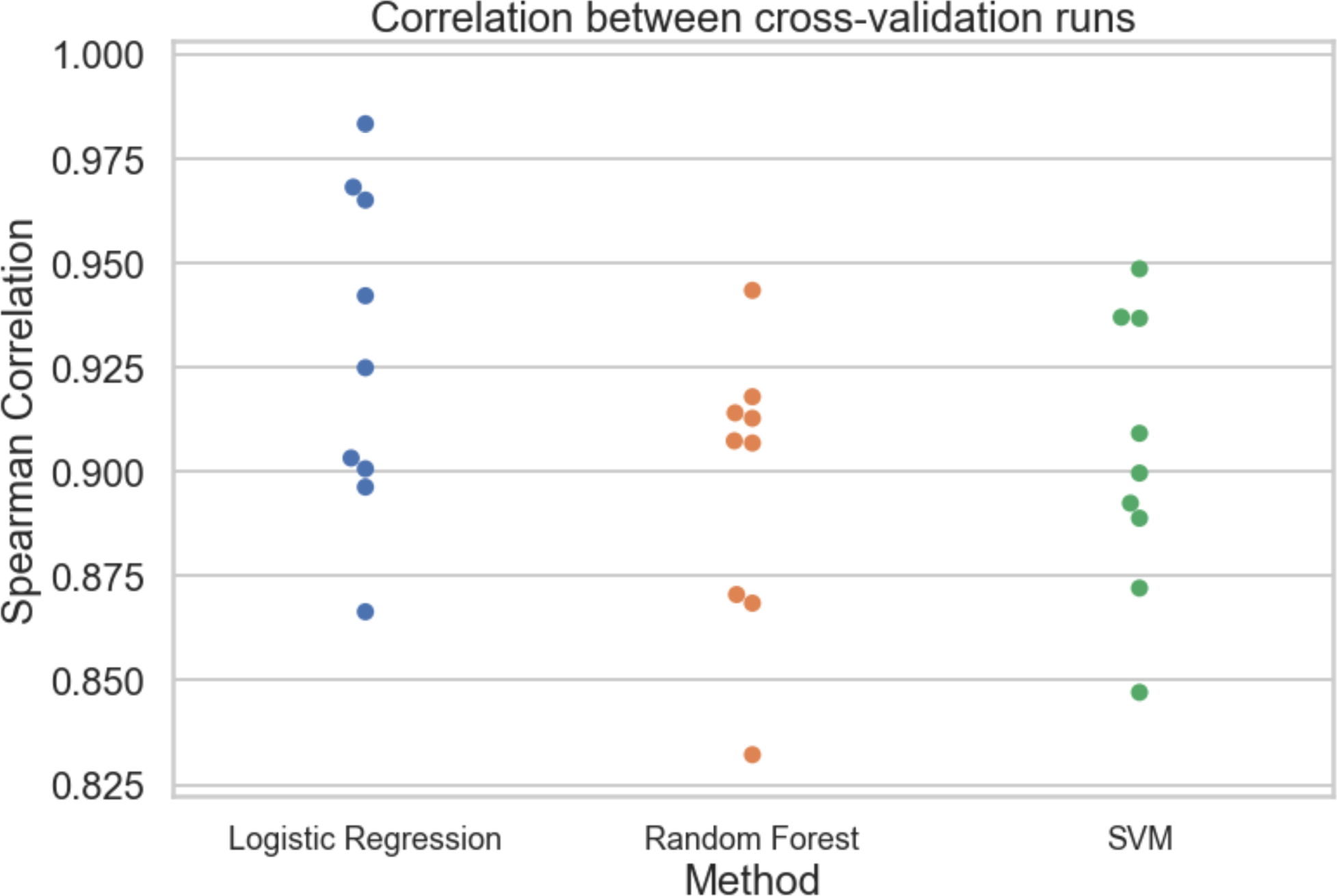
Feature correlation between cross-validation for the real data.

**Supplemental Table 1.** Spearman Correlation between variable importance scores (Random Forests) and model coefficients (Support Vector Machine and Logistic Regression) for the SPRINT trial data.

|  | Random Forest |  | Support Vector Machine |  | Logistic Regression |  |
| --- | --- | --- | --- | --- | --- | --- |
|  | Correlation | P-Value | Correlation | P-Value | Correlation | P-Value |
| Real - Non-Private | 0.7207 | 7.1518e-07 | 0.5279 | 9.35794e-04 | 0.6973 | 2.2950e-06 |
| Real - Private | 0.6123 | 7.21008e-05 | 0.6517 | 1.6644e-05 | 0.6893 | 3.3310e-06 |

**Supplemental Table 2.**
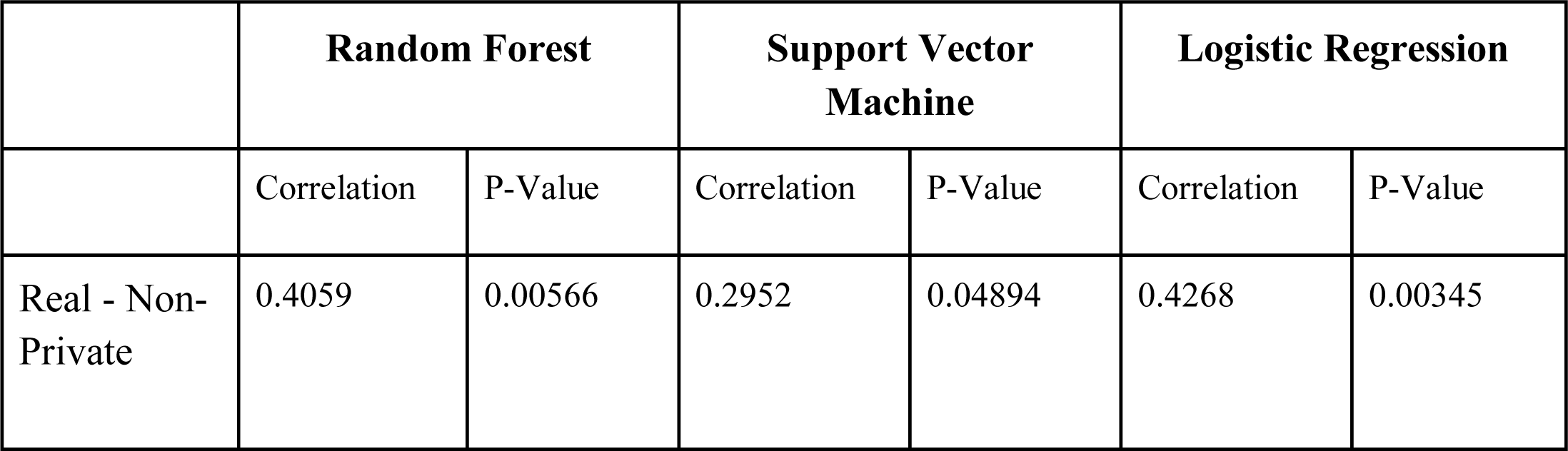
Spearman Correlation between variable importance scores (Random Forests) and model coefficients (Support Vector Machine and Logistic Regression) for the MIMIC critical care data.

|  | Random Forest |  | Support Vector Machine |  | Logistic Regression |  |
| --- | --- | --- | --- | --- | --- | --- |
|  | Correlation | P-Value | Correlation | P-Value | Correlation | P-Value |
| Real - Non-Private | 0.4059 | 0.00566 | 0.2952 | 0.04894 | 0.4268 | 0.00345 |

